# Comprehensive Characterization of Endogenous Phospholamban Proteoforms Enabled by Photocleavable Surfactant and Top-down Proteomics

**DOI:** 10.1101/2023.04.12.536120

**Authors:** Holden T. Rogers, David S. Roberts, Eli J. Larson, Jake A. Melby, Kalina J. Rossler, Austin V. Carr, Kyle A. Brown, Ying Ge

**Author notes:** To whom correspondence may be addressed: Ying Ge, 1111 Highland Ave., WIMR II 8551, Madison, WI 53705.

## Abstract

Top-down mass spectrometry (MS)-based proteomics has become a powerful tool for analyzing intact proteins and their associated post-translational modification (PTMs). In particular, membrane proteins play critical roles in cellular functions and represent the largest class of drug targets. However, the top-down MS characterization of endogenous membrane proteins remains challenging, mainly due to their intrinsic hydrophobicity and low abundance. Phospholamban (PLN) is a regulatory membrane protein located in the sarcoplasmic reticulum and is essential for regulating cardiac muscle contraction. PLN has diverse combinatorial PTMs and their dynamic regulation has significant influence on cardiac contractility and disease. Herein, we have developed a rapid and robust top-down proteomics method enabled by a photocleavable anionic surfactant, Azo, for the extraction and comprehensive characterization of endogenous PLN from cardiac tissue. We employed a two-pronged top-down MS approach using an online reversed-phase liquid chromatography tandem MS (LC-MS/MS) method on a quadrupole time-of-flight (Q-TOF) MS and a direct infusion method via an ultrahigh-resolution Fourier-transform ion cyclotron resonance (FTICR) MS. We have comprehensively characterized the sequence and combinatorial PTMs of endogenous human cardiac PLN. We have shown the site-specific localization of phosphorylation to Ser16 and Thr17 by MS/MS for the first time and the localization of S-palmitoylation to Cys36. Taken together, we have developed a streamlined top-down targeted proteomics method for comprehensive characterization of combinatorial PTMs in PLN toward better understanding the role of PLN in cardiac contractility.

## Introduction

Membrane proteins play critical roles in a wide variety of cellular functions and represent the largest class of drug targets.^1–3^ The protein activity is modulated through protein-protein interactions and post-translational modifications (PTM).^4–6^ Alterations in the PTM state of a membrane protein can have a significant impact on protein structure and function, which can lead to the development of disease.^5,7^ Despite their significance, membrane proteins are severely underrepresented in proteomic analysis as characterization from endogenous sample remains difficult.^8^ Challenges including hydrophobicity, low expression levels, as well as the overall complexity of the human proteome make the extraction, purification, and characterization of membrane proteins extremely difficult.^4,8,9^

Phospholamban (PLN) is a transmembrane protein localized in the sarcoplasmic reticulum with crucial roles in calcium (Ca^2+^) handling and heart contractility.^10–16^ Importantly, the function of PLN is dynamically regulated by its PTM state and the dysregulation of PLN is linked to cardiovascular diseases.^10,17–23^ PLN is known to be post-translationally modified by phosphorylation and S-palmitoylation.^10,18^ Two phosphorylations on Ser16 and Thr17, regulated by protein kinase A and Ca^2+^/calmodulin-dependent protein kinase II respectively, are key contributors to PLN’s regulatory impact on cardiac contraction.^10–14,21,22,24^ S-palmitoylation on Cys36 by an acyltransferase highlights PTM crosstalk within PLN, as this modification has been implicated both in the enhancement of phosphorylation on Ser16 and pentamer formation once PLN is phosphorylated.^18^ The ability to investigate the presence or absence of PLN modifications directly from human cardiac tissue will be crucial to better understanding PLN’s role in disease. Therefore, it is essential to develop an analytical method capable of comprehensively characterizing PLN’s combinatorial PTMs.

Top-down mass spectrometry (MS)-based proteomics is uniquely suitable for the analysis of combinatorial PTMs.^25,26^ Unlike conventional bottom-up proteomic approach where proteins are enzymatically digested, top-down MS analyzes intact protein without digestion allowing the characterization of “proteoforms”^27^ that arise from PTMs, alternative splicing, or genetic variation.^28–31^ Furthermore, top-down MS can provide a “bird’s-eye” view to reveal different proteoforms and their relative abundance, followed by tandem MS (MS/MS) to locate the PTM sites.^25,32,33^ Despite progression in the field, top-down proteomics of membrane proteins is still difficult due to their hydrophobic nature and instability in solution.^1,4,34–38^ There is still a need for a reproducible and robust top-down proteomics method to characterize PLN due to challenges in extraction efficiency, effective chromatography, and the site-specific localization of individual PTMs.

We have previously identified a photocleavable MS-compatible surfactant (Azo), which enabled the effective solubilization of membrane proteins for top-down proteomics.^39^ Herein, we have developed Azo-enabled^39,40^ extraction method that facilitates the extraction and characterization of endogenous human cardiac PLN. We implemented a two-pronged top-down proteomics approach employing online reversed-phase liquid chromatography tandem mass spectrometry (RPLC-MS/MS) on a quadrupole time-of-flight (Q-TOF) MS and direct infusion into an ultrahigh-resolution Fourier-transform ion cyclotron resonance (FTICR) MS (**Scheme 1**). The highly reproducible and robust top-down proteomics platform resulted in the simultaneous identification and fragmentation of various PLN proteoforms for comprehensive characterization of combinatorial PTMs. This novel top-down MS-based approach enabled comprehensive PTM mapping of PLN, including localization of S-palmitoylation to Cys36 and site-specific localization of phosphorylation to Ser16 and Thr17 by MS/MS for the first time.

### Experimental Section

#### Chemicals and reagents

All reagents were purchased from Millipore Sigma (Burlington, MA, USA) unless otherwise noted. Buffers were prepared with LC-MS-grade water from Milli-Q (Millipore, Corp., Billerica, MA, USA). Isopropanol (IPA) and acetonitrile (ACN) were purchased from Fisher Scientific (Fair Lawn, NJ, USA).

#### Human Cardiac Tissue Collection

Heart tissue was obtained from the University of Wisconsin (UW)-Madison Organ and Tissue Donation. The procedures for the collection of human donor heart were approved by the Institutional Review Board (IRB) of the UW-Madison. Available deidentified clinical data for the heart tissue used in this study can be found in **Table S1**.

#### Protein Extraction

Proteins were extracted from ∼15 mg of cryopulverized human donor cardiac tissue using a two-stage extraction at 4 ºC. Cytosolic proteins were depleted twice with 150 μL ammonium bicarbonate (ABC) buffer (25 mM ABC pH = 8.0, 10 mM L-Methionine, 5 mM TCEP, 1 mM PMSF, 2.5 mM EDTA, 1x Protease and Phosphatase Inhibitor Cocktail). The homogenate was centrifuged at 21,000 *g* for 30 min at 4 ºC after which the supernatant was discarded. After centrifugation, the pellet was re-homogenized in 150 μL Azo buffer (25 mM ABC pH = 8.0, 0.4% Azo in 25 mM ABC, 10 mM L-Methionine, 5 mM TCEP, 1 mM PMSF, 2.5 mM EDTA, 1x Protease and Phosphatase Inhibitor Cocktail) to solubilize membrane proteins, including PLN. The homogenate was centrifuged at 21,000 *g* for 30 min at 4 ºC and the supernatant was collected. The supernatant was centrifuged again at 21,000 *g* for 30 min at 4 ºC to remove any other cell debris.

#### Sample Preparation

Extracted proteins were buffer exchanged using a methanol/chloroform/water precipitation and resolubilization adapted from previous methods.^41,42^ 340 μL of cold LC/MS-grade water (4 ºC) was added to 60 μL of protein solution. 400 μL of cold methanol (−20 ºC) was then added to the protein solution and vortexed for 30 s followed by 100 μL of cold chloroform (−20 ºC) and an additional 30 s of vortexing. The sample was centrifuged for 10 min at 18,000 *g* at 4°C after which a biphasic mixture was created with a protein pellet present at the interface. The top layer of the solution was discarded without disturbing the protein pellet. 400 μL of cold methanol (−20 ºC) was then added to the sample and gently vortexed. The sample was centrifuged for 10 min at 18,000 *g* at 4° C after which the supernatant was discarded. The previous two steps were repeated two additional times. Protein pellets were resolubilized with a small volume of 80% formic acid (−20 ºC) and diluted to 5% formic acid with 10:10:80 IPA:ACN:water. Samples were UV-irradiated for approximately 3 min to degrade of any remaining Azo not removed by buffer exchange. Concentrations of samples were determined via Bradford assay and protein injection amounts were normalized prior to LC-MS/MS analysis.

#### Online Top-down LC-MS/MS Data Acquisition

Top-down LC-MS/MS was carried out using a NanoAcquity ultra-performance liquid chromatography (UPLC) system or Acquity UPLC M-Class system (Waters) coupled to a high-resolution maXis II Q-TOF-MS or Impact II Q-TOF-MS (Bruker Daltonics). Samples were injected onto a home-packed reversed phase column (C2, 150 × 0.5 mm, 5-μm particle size, 300 Åpore size) using a gradient of 10 to 90% mobile phase B (0-5 min-10% B, 5-45 min-15-65% B, 45-50 min-65-90% B, 50-55 min-90% B, 55.1-60 min-10% B; mobile phase A: 0.2% formic acid in water; mobile phase B: 0.2% formic acid in 50:50 ACN:IPA). Flow rate was set to 20 μL/min with a column temperature of 60 ºC. Mass spectra were taken at a scan rate of 0.5 Hz over 550-2000 *m/z* range. Specific MS parameters can be found in **Supplemental Note 1**. For targeted collision-induced dissociation (CID) MS/MS experiments, the precursor ion was set with a width of at least 6 *m/z*. Collision energies were set to values ranging from 15-50 eV depending on the targeted proteoform.

#### Direct Infusion Top-down MS Data Acquisition

Desalted and UV-irradiated protein extract (1.79 mg/mL) was directly infused into a Bruker 12 T SolariX FTICR mass spectrometer using an Advion TriVersa NanoMate electrospray source. Gas pressure and spray voltage was set between 0.25-0.60 psi and 1.3-1.6 kV, respectively. For MS and MS/MS analysis, optimal values for Funnel 1 RF, Skimmer 1, Collision Energy, Nebulizer, Dry Gas, Dry Temp, Acquisition Size, Accumulation Time, and Transient Length were determined to be 150 V, 50 V, 4 eV, 0.1 Bar, 3.0 L/min, 180 ºC, 1 to 2 M, 5.0 to 10.0 s, and 1.15 to 2.31 s, respectively. Additional parameters including Capillary Exit, Deflector plate, Octopole Frequency, Octopole RF Amplitude, Collision Cell RF Frequency and Collision Cell RF Amplitude were set to 240 V, 220 V, 1.0, 600 Vpp, 1.0, and 2000 Vpp, respectively. A minimum of 50 scans were accumulated for each sample with a resolution of 530,000 at 400 *m/*z. Prior to MS/MS, the precursor ion isolation was set to a width of 2-4 *m/z*. Collision was set to energies ranging from 20-30 eV. FTICR mass spectra were calibrated externally using 50 mM cesium iodide in water with a 200-5000 *m/z* range.

#### Data Analysis

DataAnalysis (Version 4.3; Bruker Daltonics) software was used to process and analyze the LC-MS raw files. To quantify protein expression across samples, the Top 3 most abundant charge states’ ions (most abundant ± 0.5 *m/z*) of all major proteoforms and their first oxidation state were retrieved collectively as one extracted peak in the extracted ion chromatogram (EIC). The area under curve (AUC) was manually determined for each proteoform using DataAnalysis. To quantify protein modifications, the relative abundances of specific modifications (mono-/bis-phosphorylated and palmitoylated bis-phosphorylated) were calculated from their respective AUC values and reported as corresponding ratios across three extraction replicates. Monoisotopic masses from MS and MS/MS spectra are reported. Tandem mass spectra were exported from DataAnalysis software and analyzed using MASH Native (Version 1.1).^43,44^ eTHRASH was used to deconvolute exported files using default parameters. An error tolerance of 20 ppm and 5 ppm was used for fragments produced from the online LC-MS/MS and direct infusion methods, respectively. The minimum fragment size accepted was three amino acids. Internal fragments are labeled using the format *by* [start amino acid] - [end amino acid] [charge state].

## Results and Discussion

### Online LC-MS Analysis of PLN Proteoforms

In this study, we sought to develop a simple, robust, and highly reproducible top-down LC-MS/MS method for the comprehensive characterization of PLN from endogenous tissue samples. To address the challenges in sample preparation we used a photocleavable MS-compatible surfactant, Azo,^39^ for effective solubilization and extraction of PLN from heart tissue. Depletion of cytosolic proteins before the addition of Azo served to reduce the proteome complexity prior to MS analysis. An important criterion for the online LC-MS/MS approach was to produce a high-throughput method that enabled the simultaneous analysis of PLN’s combinatorial PTM states in a single run. To accomplish this, a strong emphasis was placed on front-end RPLC separation development. To maximize throughput, we initially surveyed a condensed 28-min gradient, which demonstrated high reproducibility (**Figure S1A and S1B**) and high-quality fragmentation of phosphorylated PLN proteoforms. However, significant coelution of non-specific proteins and peptides across key *m/z* windows corresponding to PLN (**Figure S1C**) prevented the detection of palmitoylated PLN proteoforms.

To increase separation efficiency, an extended 60-min gradient was then implemented. Importantly, the 60-min gradient resulted in the detection of both phosphorylated and palmitoylated PLN proteoforms. Representative total ion chromatograms (TIC) and extracted ion chromatograms (EIC) corresponding to PLN display the elution profile of the protein extraction (**Figure 1A and 1B**). Across three extraction replicates, EICs for the mono-phosphorylated, bis-phosphorylated, and palmitoylated bis-phosphorylated proteoforms demonstrated high reproducibility (**Figure S2A**). Mass spectra normalized for intensity (**Figure S2B)** and AUC analysis of individual PLN proteoforms (**Figure S2C**) showed comparable relative abundances. PLN proteoforms also displayed strong linear responses upon increasing injection amounts (**Figure S3**). Mass spectra averaged across the elution window of individual PLN PTM states demonstrated minimal coelution (**Figure 1C-D**). With the extended gradient, we concurrently examined five unique PLN proteoforms that contained different combinations of acetylation, phosphorylation, and palmitoylation **(Table S2)**. Although PLN was generally observed eluting at a high organic mobile-phase percentage (>65% B), not all proteoforms eluted at the same time. Given the small size (6.1 kDa) and physiochemical properties of PLN, PTMs such as S-palmitoylation can have a significant impact on the PLN’s retention within the column. This is highlighted by a peak-to-peak retention shift of 3.2-min between PLN’s palmitoylated and non-palmitoylated forms. The aforementioned separation proved to be desirable for MS/MS characterization as it enabled a targeted fragmentation approach for individual proteoforms.

**Figure 1.**
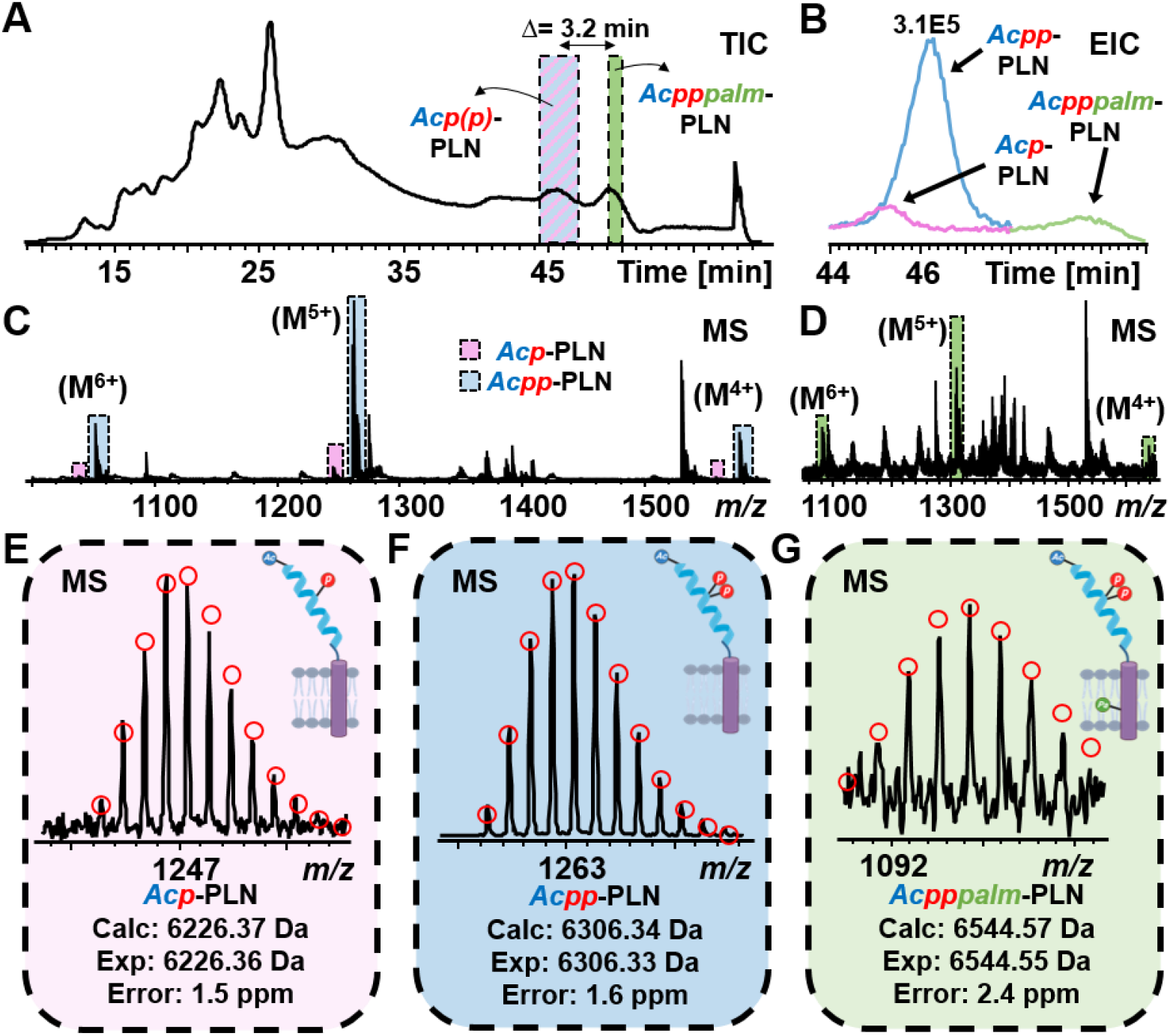
Online LC-MS analysis of PLN proteoforms. **(A)** Representative TIC exhibiting the retention differences of mono- and bis-phosphorylated PLN proteoforms compared to the palmitoylated bis-phosphorylated PLN proteoform. The change in reported elution time is determined by peak-to-peak difference (3.2 min). **(B)** Representative EIC exhibiting the retention shift and relative abundance differences of PLN proteoforms. **(C)** MS of mono- (pink) and bis-phosphorylated (blue) PLN proteoforms across *m/z* space. **(D)** MS of palmitoylated bis-phosphorylated (green) PLN proteoforms across *m/z* space. **(E-G)** Zoomed-in MS spectra of mono-phosphorylated **(E)**, bis-phosphorylated **(F)**, and palmitoylated bis-phosphorylated **(G)** PLN proteoforms. Data collected using Impact II. Theoretical isotopic fits are shown in red circles. Monoisotopic masses are reported.

The mono-phosphorylated (**Figure 1E**) and bis-phosphorylated (**Figure 1F**) proteoforms were the two most abundant PLN proteoforms identified from MS-level analysis. The experimentally determined mono-isotopic masses (Exp) for both proteoforms were within 1.6 ppm mass accuracy of theoretically calculated masses (Calc). Although eluting later in the gradient, the palmitoylated bis-phosphorylated proteoform (**Figure 1G**), was identified with high mass accuracy as the experimentally determined mono-isotopic mass had sub-2.5 ppm mass error. The acetylated PLN proteoform and the palmitoylated mono-phosphorylated PLN proteoform were also identified in this donor human heart tissue (**Figure S4**); however, they were lower in abundance compared to the mono- and bis-phosphorylated proteoforms. Regardless of the specific proteoform or their relative abundance to each other, the M^5+^ charge state was consistently the most abundant throughout all experiments. Overall, the top-down proteomics method provides a “bird’s-eye” view of PLN’s combinatorial PTMs with high accuracy mass measurements and high reproducibility.

### Localization of PLN’s Palmitoylation Site

Following intact MS analysis, further confirmation of PLN’s proteoforms was achieved by MS/MS. We first looked at the palmitoylated bis-phosphorylated PLN proteoform. In membrane proteins, palmitoylation has significant roles impacting protein conformation, association to membrane domains, protein-protein interactions and PTM crosstalk.^45^ S-palmitoylation, a 15-hydrocarbon chain added on the Cys36 residue of PLN, significantly increases the hydrophobicity of PLN and is known to augment phosphorylation and contribute to pentamer formation.^18^ However, S-palmitoylation is a highly labile PTM that has the potential of being removed if significant collision energy is applied.^46^ Therefore, to characterize the palmitoylated proteoform, a combination of lower collision energies (15, 20, 25, 30 eV) were evaluated to minimize the removal of the labile PTM during analysis (**Figure 2**). After isolation and fragmentation of the palmitoylated PLN proteoform, validated fragments resulted in full sequence coverage and 25% bond cleavage (20 fragment ions: 13 *b*-ions/6 *y*-ions). Fragmentation results of the individual collision energies along with annotated MS/MS spectra can be found in the supplemental info (**Figure S5)**. Together, fragment ions *b*_*38*_ ^*4+*^, *y*_*7*_ ^*1+*^ and *y*_*12*_ ^*1+*^ demonstrate site-specific localization of S-palmitoylation to Cys36 rather than Cys41 or Cys46. Additionally, there was no evidence of significant removal of palmitoylation upon fragmentation as demonstrated by the negligible presence of intact bis-phosphorylated PLN (**Figure S6**) and the absence of *b/y*-ion fragments corresponding to any non-palmitoylated proteoform. Electron-transfer dissociation (ETD) and electron-capture dissociation (ECD) are two fragmentation techniques known to also preserve labile modifications.^46,47^ For this reason, both ETD and ECD were attempted to provide complementary fragmentation of PLN. However, in the case of PLN, MS/MS efforts with ECD and ETD did not produce robust fragmentation.

**Figure 2.**
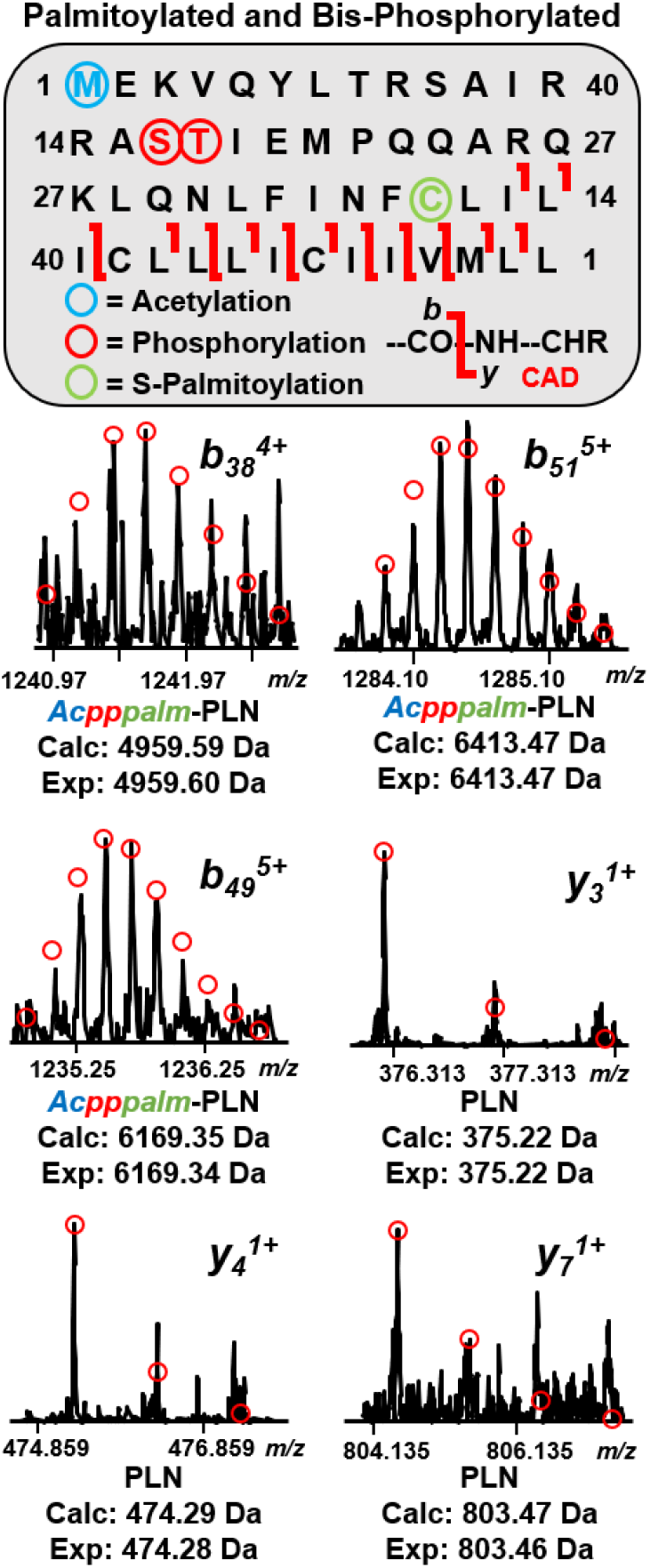
Top-down MS of bis-phosphorylated palmitoylated PLN proteoform. Representative fragment ion spectra of bis-phosphorylated palmitoylated PLN (*b*_*38*_^*4+*^, *b*_*51*_^*5+*^, *b*_*49*_^*5+*^, *y*_*3*_^*1+*^, *y*_*4*_^*1+*^, *y*_*7*_^*1+*^). The percentage of bond cleavages is 25%. Sequence table reports combined fragmentation coverage of PLN proteoforms across four different collisional energies (15, 20, 25, and 30 eV). Data collected using Impact II. Theoretical isotopic fits are created using MASH Native v. 1.1. Monoisotopic masses are reported.

### Localization of PLN’s Phosphorylation Sites

Previous proteomic studies employing offline prefractionation techniques were unable to distinguish site occupancy of phosphorylated proteoforms to the residues of Ser16 and Thr17 by MS/MS.^48^To achieve this, we started with an online LC-MS/MS analysis for the mono- and bis-phosphorylated PLN proteoform (**Figure 3**). The M^5+^ charge states of the mono-and bis-phosphorylated PLN proteoforms were isolated and fragmented by CID in the online RPLC-MS/MS method (**Figure 3A and 3B**). Four different collisional energies (20, 30, 40, 50 eV) were used to achieve high bond cleavages and their spectra were combined. Full sequence coverage was achieved for both proteoforms. For the mono-phosphorylated proteoform (**Figure 3A**), MS/MS fragmentation produced 54% bond cleavage (40 fragment ions: 28 *b*-ions/12 *y*-ions). For the bis-phosphorylated proteoform (**Figure 3B**), MS/MS fragmentation produced 65% bond cleavage (47 fragment ions: 34 *b*-ions/13 *y*-ions). The differences in bond cleavage can be attributed to the relative intensity of each proteoform, as the bis-phosphorylated PLN proteoform was significantly higher in abundance compared to its mono-phosphorylated counterpart. MS/MS results of the individual collision energies and annotated spectra can be found in the supplemental info (**Figure S7 and Figure S8)**.

**Figure 3.**
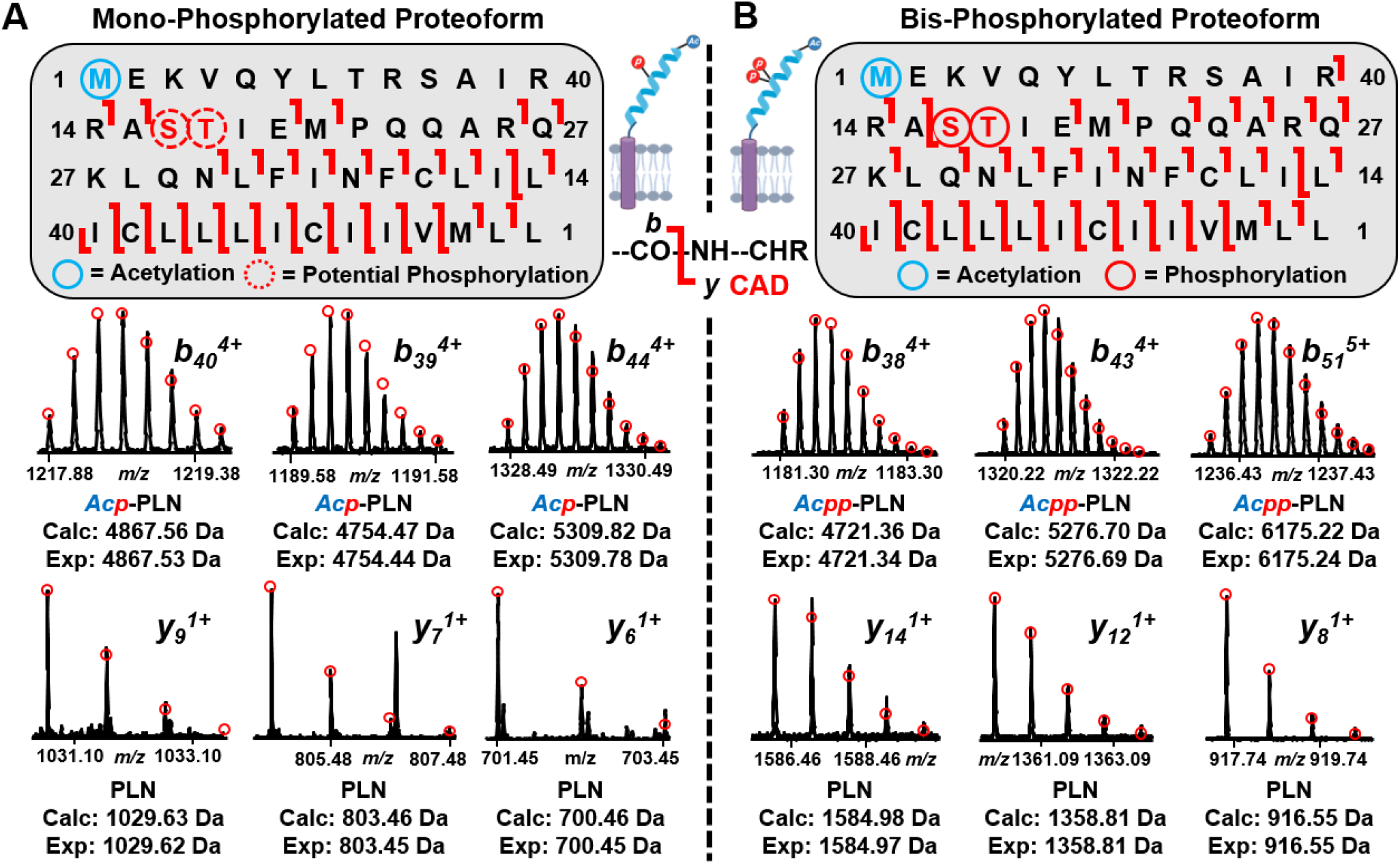
Online RPLC-MS/MS analysis of mono- and bis-phosphorylated PLN proteoforms. **(A)** Representative fragment ion spectra of mono-phosphorylated PLN (*b*_*40*_^*4+*^, *b*_*39*_^*4+*^, *b*_*44*_^*4+*^, *y*_*9*_^*1+*^, *y*_*7*_^*1+*^, *y*_*6*_^*1+*^). The percentage of bond cleavages is 54%. **(B)** Representative fragment ion spectra of bis-phosphorylated PLN (*b*_*38*_^*4+*^,*b*_*43*_^*4+*^,*b*_*51*_^*5+*^,*y*_*14*_^*1+*^,*y*_*12*_^*1+*^,*y*_*8*_^*1+*^).The percentage of bond cleavages is 65%. Sequence tables report combined fragmentation coverage of PLN proteoforms across four different collisional energies (20, 30, 40, and 50 eV). Data collected using Impact II and maXis II. Theoretical isotopic fits are created using MASH Native v. 1.1. Monoisotopic masses are reported.

With an elution window spanning 1 to 2 min, our online LC-MS/MS method was able to achieve sufficient fragmentation for regional localization for PLN’s phosphorylations. For mono-phosphorylated PLN, fragment ions *b*_*15*_^*2+*^ and *b*_*19*_ ^*2+*^ narrowed the region of the single phosphorylation to a residue between Ser16 and Glu19 (**Figure 3A)**. Likewise, for bis-phosphorylated PLN, fragment ions *b*_*15*_ ^*2+*^, *b*_*19*_ ^*2+*^, and *y*_*37*_ ^*3+*^ support the presence of two phosphorylations to the same Ser16 to Glu19 region (**Figure 3B)**. The data collected demonstrated the overall effectiveness of our online MS/MS method in fragmenting the mono- and bis-phosphorylated proteoforms, as it resulted in the highest reported bond cleavage for PLN. Moreover, the online-LC-MS/MS approach developed here presents a rapid, straightforward, and robust platform that enables the simultaneous characterization of PLN proteoforms with high sequence coverage. However, site-specific occupancy of phosphorylated proteoforms to the known residues of Ser16 and Thr17 was not yet distinguished.

To achieve site-specific localization, we transitioned to a second method where we directly infused complex protein extract into a Bruker SolariX 12 T FTICR MS (**Figure 4A**). The FTICR MS allowed the accumulation of PLN over an extended period, potentially enabling additional fragmentation to complement our online LC-MS/MS method. Although there was no separation prior to direct infusion, the high-resolution capability of this instrument enabled clean isolation of an *m/z* window that corresponded to the bis-phosphorylated PLN proteoform with one oxidation (**Figure 4B**). Following successful isolation of the bis-phosphorylated precursor with sub-1.0 ppm MS mass error, fragmentation was accomplished with CID where two key internal fragments were collected. The first fragment (**Figure 4C**), denoted as *by* _(17-43)_^2+^, is a 27-residue fragment that that contains one phosphorylation PTM. The second fragment (**Figure 4D**), denoted as *by*_(10-35)_ ^2+^, is a 26-residue fragment that contains two phosphorylation PTMs. The two internal fragments above, in combination with N/C-terminal fragments (*b*_*13*_^*2+*^, *b*_*14*_^*2+*^, *b*_*15*_^*2+*^, *y*_*37*_^*3+*^) collected from our online RPLC-MS/MS experiments (**Figure S9**), demonstrate site-specific localization of phosphorylation to Ser16 and Thr17. The use of internal fragments in top-down proteomic analysis has previously been ignored due to specific challenges in the assignment of fragment spectra.^49–51^ As a protein increases in size, there is an exponential increase in the number of internal fragments that can be produced, complicating analysis with false discovery rates.^49^ To verify that the spectra collected corresponded to the specific internal fragments for phosphorylation localization, we generated a complete list of possible internal and terminal fragment ions with MASH Native (**Supplemental Spreadsheet**) and we illustrated the overlap of our experimentally obtained internal fragment ions with theoretical ion distributions. In the list of theoretical fragments, there was no other terminal or internal fragment with a monoisotopic mass corresponding to the two experimentally obtained masses, further supporting the assignment of *by*_(10-35)_ ^2+^and *by*_(17-43)_ ^2+^ and localization to Ser16 and Thr17.

**Figure 4.**
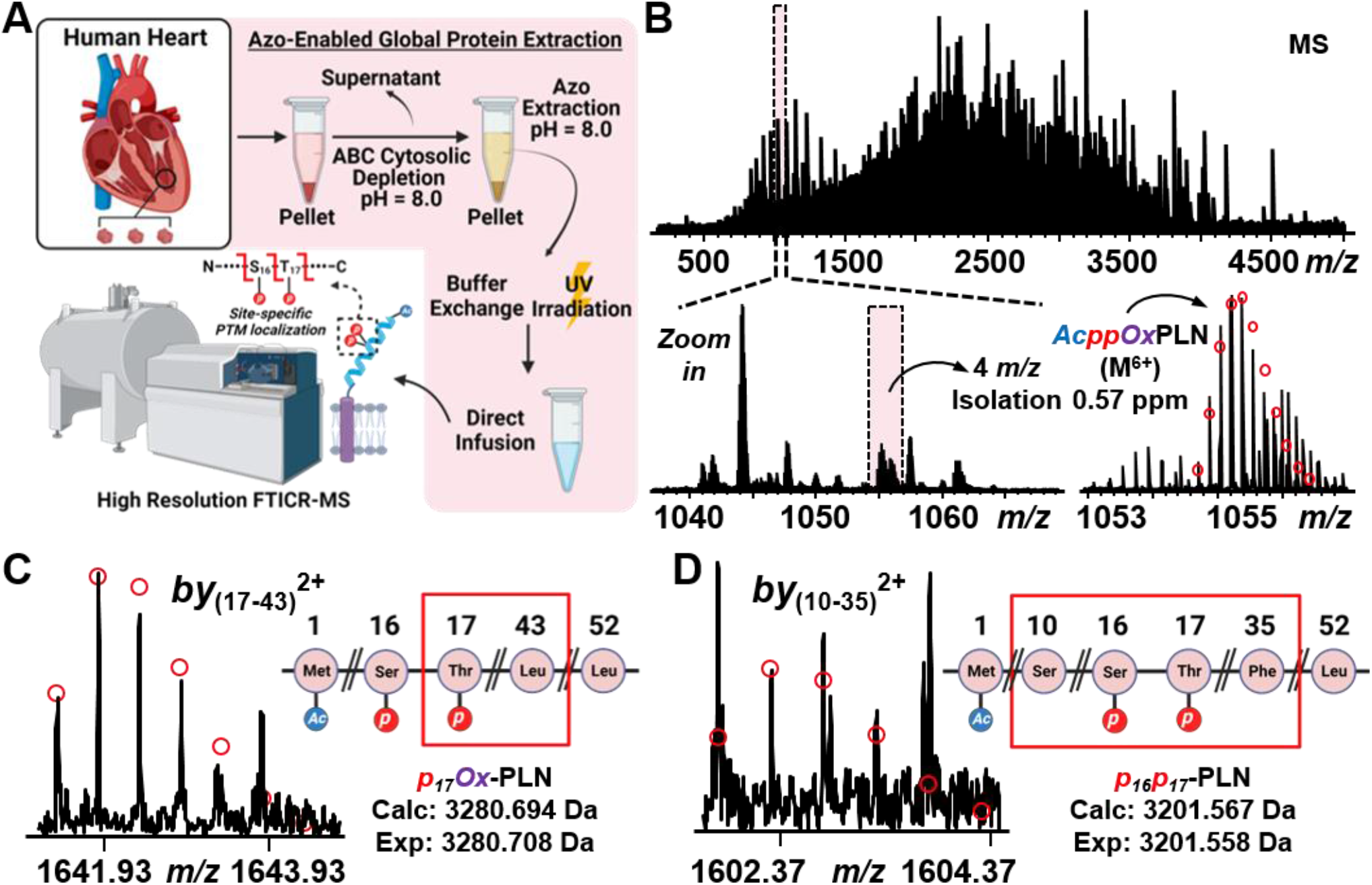
PLN site-specific phosphorylation localization. **(A)**. Schematic for direct infusion onto FTICR MS. Proteins were extracted from ∼15 mg of human donor cardiac tissue using a two-stage extraction previously described. Protein extract (1.79 mg/mL) was directly infused on a Bruker SolariX 12 T FTICR MS for analysis. **(B)** Representative MS spectrum of entire *m/z* range (500-5000 m/z). Zoomed-in window shows MS spectra after a 20 *m/z* and 4 *m/z* isolation centered on 1055 *m/z*, which represents the N-terminally acetylated and bis-phosphorylated PLN proteoform with one oxidation. **(C-D)** MS/MS analysis of isolated peak using CAD (28 eV). Representative fragment ion spectra of internal fragments (*by*_(17-43)_^2+^ and *by*_(10-35)_^2+^), in conjunction with other N/C-terminal fragments, demonstrate site specific localization of each phosphorylation, one on Thr17 and the other on Ser16. Theoretical isotopic fits are created using MASH Native v. 1.1. Monoisotopic masses are reported.

Notably, this is the first MS-based approach to enable precise localization of phosphorylation sites in PLN from human cardiac tissue. This result is critical for the endogenous characterization of PLN as prior *in vitro* studies have shown that Ser10 and Thr8 are potential phosphorylation sites for PLN.^14,21,52^ However, oftentimes there are limitations with studying PTMs through *in vitro* studies, as there can be modifications that might not recapitulate endogenous molecular forms. The fragment ions here provide an unambiguous location of PLN’s phosphorylation sites directly from human heart tissue which will be important for future studies that attempt to distinguish enzymatic pathways and the mechanisms of cardiac contractility.

## Conclusion

In summary, we have developed a novel targeted MS-based top-down proteomics method to characterize endogenous human cardiac PLN with a photocleavable anionic surfactant, Azo. We employed a two-pronged approach for the systematic analysis of PLN, one utilizing an online RPLC-MS/MS method on a Q-TOF-MS and the other exploiting direct infusion via an ultrahigh-resolution FTICR MS. For the first time, we have unambiguously localized PLN phosphorylation to Ser16 and Thr17 both of which are important in the regulation of cardiac contraction. Moreover, we have also localized S-palmitoylation, a modification necessary for phosphorylation enhancement, to Cys36. We believe this targeted top-down MS method will be highly valuable for understanding the regulatory roles of combinatorial PTMs of PLN in cardiac contractility and diseases.

### Associated Content

Data Availability Statement: The mass spectrometry proteomics data have been deposited to the ProteomeXchange Consortium via the PRIDE partner repository with the data set identifier PXD041013 and the MassIVE repository with identifier MSV000091522.

## Supplementary Notes

Note 1. Q-TOF-MS parameters

## Supplementary Tables

Table S1. Deidentified patient information Table S2. PLN proteoforms

## Supplementary Figures

Figure S1. Reproducibility of Azo-enabled extraction method with shorter 28-minute gradient

Figure S2. Reproducibility of Azo-enabled extraction method

Figure S3. Linear injection response of online LC-MS method

Figure S4. Low abundant PLN proteoforms

Figure S5. Online RPLC-MS/MS analysis of individual collision energies for the palmitoylated bis-phosphorylated PLN proteoform

Figure S6. Impact of CAD energy on palmitoylated PLN proteoform

Figure S7. Online RPLC-MS/MS analysis of individual collision energies for the mono-phosphorylated PLN proteoform

Figure S8. Online RPLC-MS/MS analysis of individual collision energies for the bis-phosphorylated PLN proteoform

Figure S9: Fragment ions supporting phosphorylation localization

## Supplementary excel spreadsheet

Supplemental Spreadsheet. Excel sheet containing theoretical fragments of PLN

### Author Contributions

H.T.R., D.S.R., and E.J.L. designed the experiments and overall approach. H.T.R. prepared samples, collected data, analyzed the data, and wrote the manuscript. D.S.R. assisted in supervising the work, data collection, formal analysis, and revised the manuscript. E.J.L assisted in data collection. K.A.B., J.A.M., K.J.R., and A.V.C. contributed to initial project development. Y.G. directed and supervised all aspects of the work and was responsible for project administration and funding acquisition. All authors contributed to editing and approval of the final manuscript.

### Notes

The authors declare the following competing financial interest(s): The University of Wisconsin-Madison has filed a provisional patent application P180335US01, US serial number 62/682027 (7 June 2018) on the photocleavable surfactant. Y.G., and K.A.B. are named as inventors on the provisional patent application.

## Acknowledgements

This research is supported by NIH R01 HL109810-06 (Y.G.). Y.G. would like to acknowledge NIH R01 GM125085, R01 HL096971, and S10 OD018475. D.S.R. also acknowledges the support from the American Heart Association Predoctoral Fellowship Grant No. 832615/David S. Roberts/2021. J.A.M. would like to acknowledge support from the Training Program in Translational Cardiovascular Science, T32 HL007936-21. K.J.R acknowledges the National Science Foundation Graduate Research Fellowship Program under Grant No. DGE-1747503 and the Graduate School and the Office of the Vice Chancellor for Research and Graduate Education at the University of Wisconsin-Madison, funded by Wisconsin Alumni Research Foundation. Graphics in Scheme 1 and Figure 4A were created with clipart from BioRender.com.

---

**Scheme 1.**
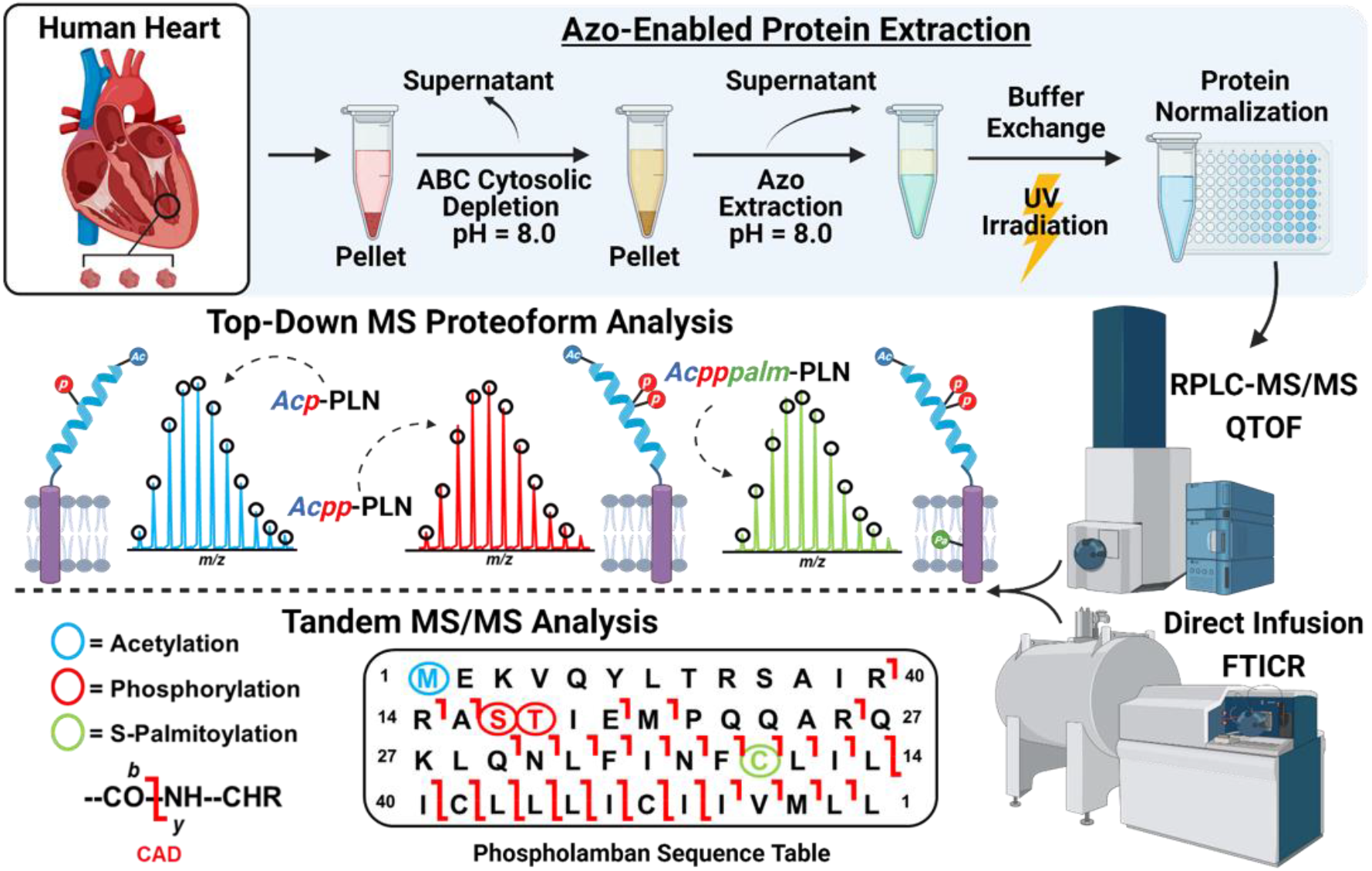
Schematic representation of top-down proteomics for comprehensive characterization of PLN proteoforms using a photocleavable surfactant, Azo. Proteins were extracted from ∼15 mg of human donor cardiac tissue using a two-stage extraction. Cytosolic proteins were depleted using an ammonium bicarbonate (ABC) buffer (pH = 8.0). After centrifugation, the pellet was homogenized in Azo buffer (pH = 8.0) to solubilize membrane proteins. Following desalting and Azo degradation, intact proteins were directly infused into a Bruker SolariX 12T FTICR MS or separated by online RPLC followed by MS analysis using a Bruker Q-TOF-MS.

