## Supplementary Information for "Comprehensive Characterization of Endogenous Phospholamban Proteoforms Enabled by Photocleavable Surfactant and Top-down Proteomics"

**Table of Contents**

Supplementary Notes

Supplementary Note 1. Q-TOF-MS parameters……….……………….……………….………S-3

Supplementary Tables

Table S1. Deidentified patient information…………………………….………...…..…….S-4

Table S2. PLN proteoforms ……………………………………………………...….S-5

Supplementary Figures

Supplementary Figure S1. Reproducibility of Azo-enabled extraction method with shorter 28-minute gradient……………………………………………………………..…...…..….S-6

Supplementary Figure S2. Reproducibility of Azo-enabled extraction method……..….S-7

Supplementary Figure S3. Linear injection response of online LC-MS method………..S-8

Supplementary Figure S4. Low abundant PLN proteoforms……………………...….…S-9

Supplementary Figure S5. Online RPLC-MS/MS analysis of individual collision energies for the palmitoylated bis-phosphorylated PLN proteoform..………………..…..……..S-10

Supplementary Figure S6. Impact of CAD energy on palmitoylated PLN proteoform…….…………………………………………......................................S-11

Supplementary Figure S7. Online RPLC-MS/MS analysis of individual collision energies for the mono-phosphorylated PLN proteoform.……………………..………..…..…S-12

Supplementary Figure S8. Online RPLC-MS/MS analysis of individual collision energies for the bis-phosphorylated PLN proteoform……………………….…………..…..…..S-13

Supplementary Figure S9. Fragment ions supporting phosphorylation localization…..S-14

**Supplementary Note 1.**

**Q-TOF-MS parameters.** For the electrospray ionization source, End Plate Offset, Capillary, Nebulizer, Dry Gas, and Dry Temp were set to 500 V, 4500 V, 0.6 Bar, 4.5 L/min and 220 ºC, respectively. For tune settings, optimal values for Funnel 1 RF, isCID Energy, Multipole RF (for maXis II), Funnel 2 RF (for Impact II), Hexapole RF (for Impact II), Ion Energy, Collision Energy, Collision RF, Transfer Time, and Pre-Pulse Storage were determined to be 350 Vpp, 10 eV, 400 Vpp, 350 Vpp, 600 Vpp, 10 eV, 4 eV, 2600 Vpp, 110 µs, and 10 µs, respectively.

**SI Table 1. Deidentified patient information.**

| Sample Code | Source | Age | Gender | Cause of Death | Medical History |
| --- | --- | --- | --- | --- | --- |
| YG54 | University of Wisconsin Hospital and Clinic | 62 | Male | - | No cardiac history; EF of 65%; hypothyroid; high cholesterol |

**SI Table 2. PLN proteoforms.** Calculated mono-isotopic *m/z*, most abundant *m/z*, intact mono-isotopic and most abundant mass listed for PLN proteoforms across three charge states ($M^{5+}$, $M^{6+}$, $M^{4+}$). *Ac* represents N-terminal acetylation. *p* represents phosphorylation. *palm* represents palmitoylation.

| Proteoform | Charge State | Mono-isotopic Calc (*m/z*) | Most abundant Calc (*m/z*) | Mono-isotopic Calc (Da) | Most abundant Calc (Da) |
|---|---|---|---|---|---|
| *Ac*-PLN | 5+ | 1230.29 | 1231.13 | 6146.40 | 6150.63 |
| *Acp*-PLN | 5+ | 1246.28 | 1247.13 | 6226.37 | 6230.60 |
| *Acpp*-PLN | 5+ | 1262.28 | 1263.12 | 6306.34 | 6310.57 |
| *Acppalm*-PLN | 5+ | 1293.93 | 1294.77 | 6464.60 | 6468.83 |
| *Acpppalm*-PLN | 5+ | 1309.92 | 1310.77 | 6544.57 | 6548.80 |
| *Ac*-PLN | 6+ | 1025.41 | 1026.11 | 6146.40 | 6150.63 |
| *Acp*-PLN | 6+ | 1038.74 | 1039.44 | 6226.37 | 6230.60 |
| *Acpp*-PLN | 6+ | 1052.06 | 1052.77 | 6306.34 | 6310.57 |
| *Acpppalm*-PLN | 6+ | 1091.77 | 1092.47 | 6544.57 | 6548.80 |
| *Ac*-PLN | 4+ | 1537.61 | 1538.67 | 6146.40 | 6150.63 |
| *Acp*-PLN | 4+ | 1557.60 | 1558.66 | 6226.37 | 6230.60 |
| *Acpp*-PLN | 4+ | 1577.59 | 1578.65 | 6306.34 | 6310.57 |
| *Acpppalm*-PLN | 4+ | 1637.15 | 1638.21 | 6544.57 | 6548.80 |

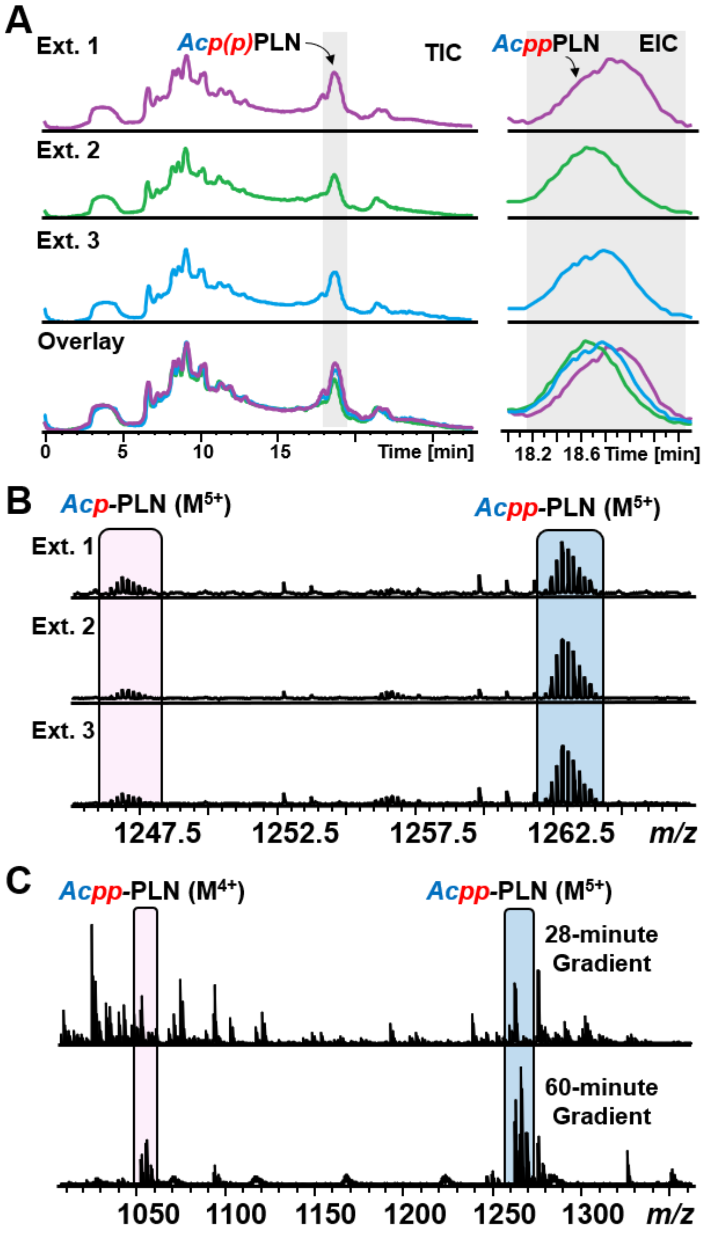


**SI Figure 1. Reproducibility of Azo-enabled extraction method with shorter 28-minute gradient. (A)** Individual and overlaid total ion chromatograms (TIC) and extracted ion chromatograms (EIC) of three extraction replicates. Flow rate was 12 µL/min. Gradient was 20 to 100% mobile phase B (0-1 min- 20% B, 1-11 min- 20-60% B, 11-15 min- 60-95% B, 15-21 min- 100% B, 21-24 min- 100% B, 24-25 min- 100-20% B, 25-28 min- 20% B). EICs are of bis-phosphorylated PLN. Grey shading indicates elution window of PLN. **(B)** MS of PLN ($M^{5+}$) proteoforms across three extraction replicates. *p* and *pp* denote mono-phosphorylated and bis-phosphorylated proteoforms, respectively. *Ac* indicates N-terminal acetylation. **(C)** MS spectra for the 28-minute and 60-minute RPLC-MS gradient demonstrating lower amounts of coelution for a longer gradient. Palmitoylated PLN was not seen in the 28-minute gradient.

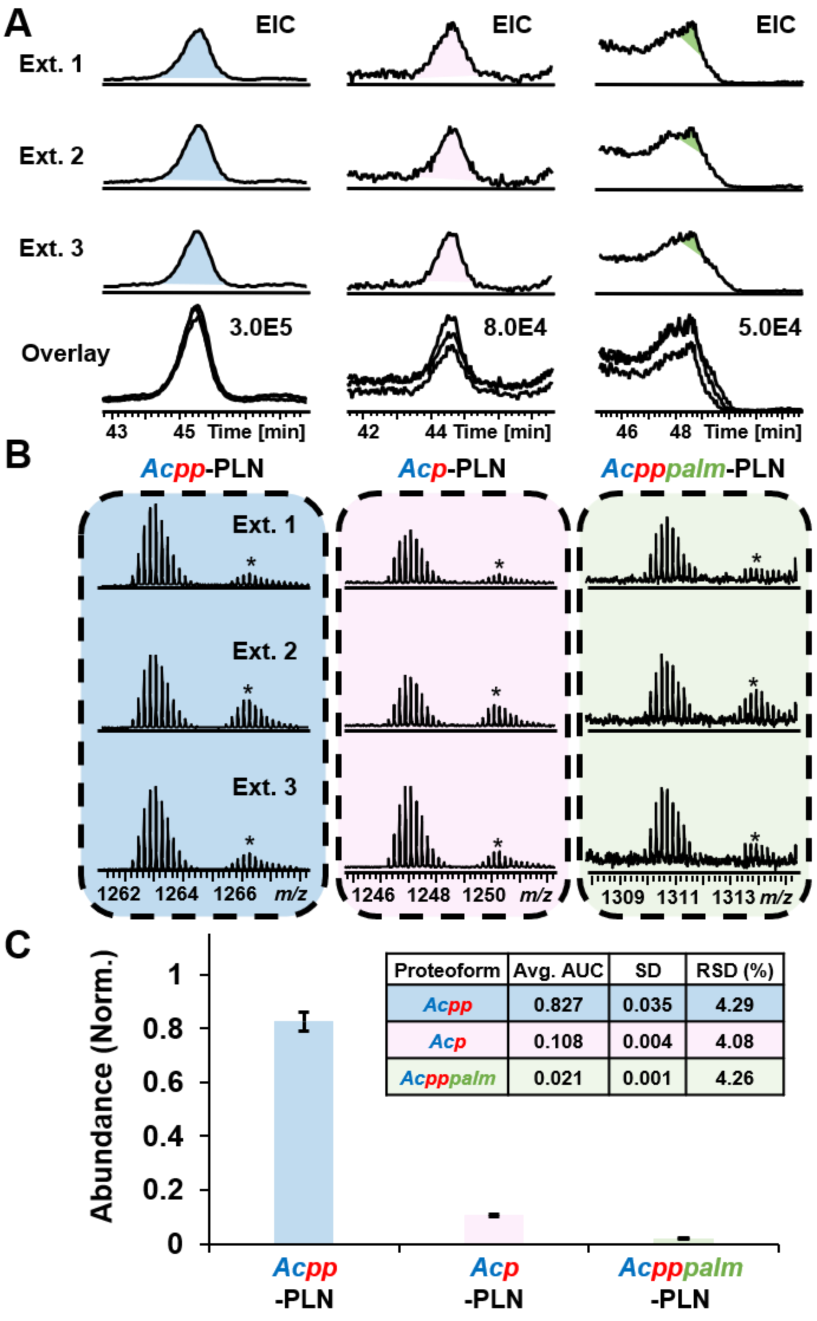


**SI Figure 2. Reproducibility of Azo-enabled extraction method. (A)** Individual and overlaid extracted ion chromatograms (EIC) of three extraction replicates for each PLN proteoform: bis-phosphorylated (blue), mono-phosphorylated (pink) and palmitoylated bis-phosphorylated (green). Shading indicates calculated AUC region. **(B)** MS of PLN ($M^{5+}$) proteoforms across three extraction replicates. *p* and *pp* denote mono-phosphorylated and bis-phosphorylated proteoforms, respectively. *Ac* indicates N-terminal acetylation. *palm* represents S-palmitoylation. The * specifies one oxidation. **(C)** AUC analysis of individual PLN proteoforms. Avg. AUC represents relative abundance of specific PLN proteoform across the three extraction replicates. Standard deviation (SD) and relative standard deviation (RSD) are provided.

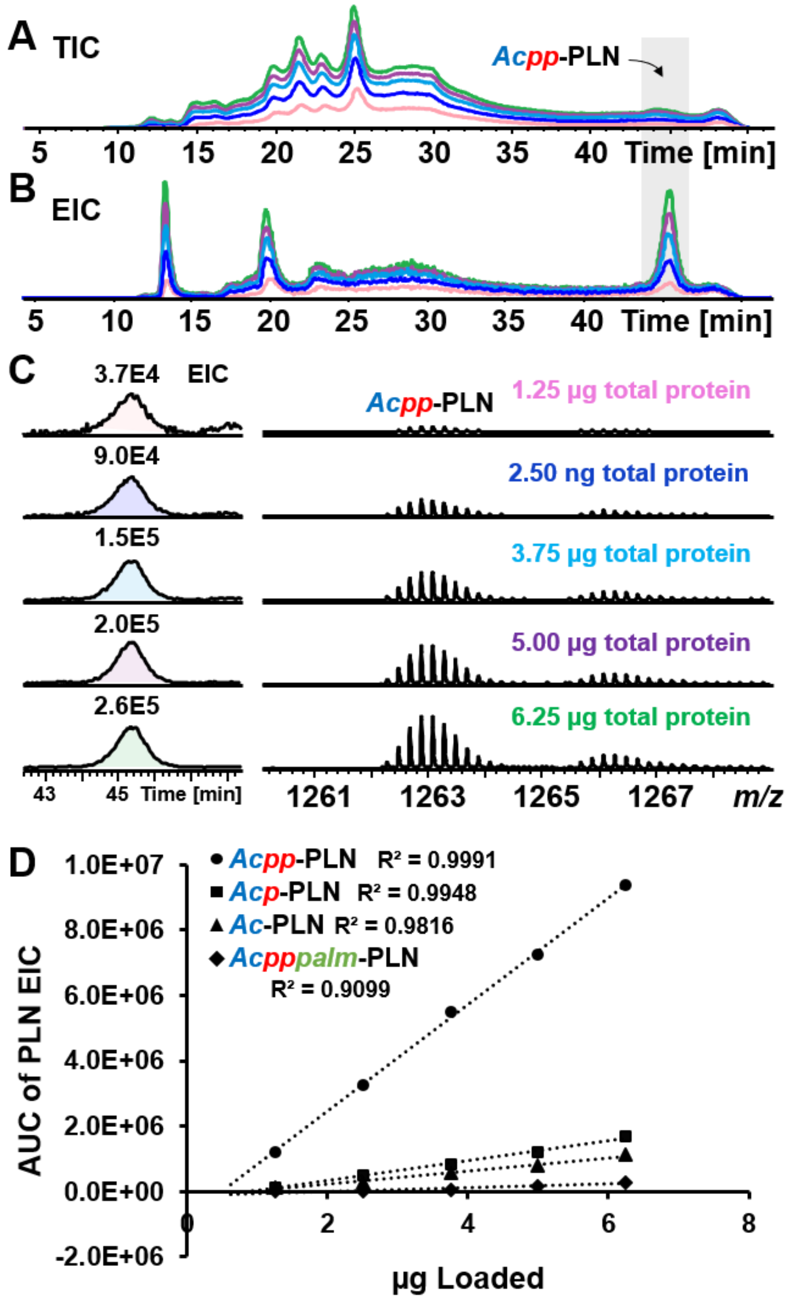


**SI Figure 3. Linear injection response of online LC-MS method. (A-B)** Representative TICs **(A)** and EICs **(B)** of five different loading amounts (1.25 µg, 2.50 µg, 3.75 µg, 5.00 µg, and 6.25 µg). PLN elution window shaded in grey. **(C)** Representative EICs and intensity normalized mass spectra of five different loading amounts (1.25 µg, 2.50 µg, 3.75 µg, 5.00 µg, and 6.25 µg). Zoomed-in EICs are of bis-phosphorylated PLN proteoform. **(D)** Linear response curve of injection amount for each individual PLN proteoform using calculated AUC. R-squared values are reported.

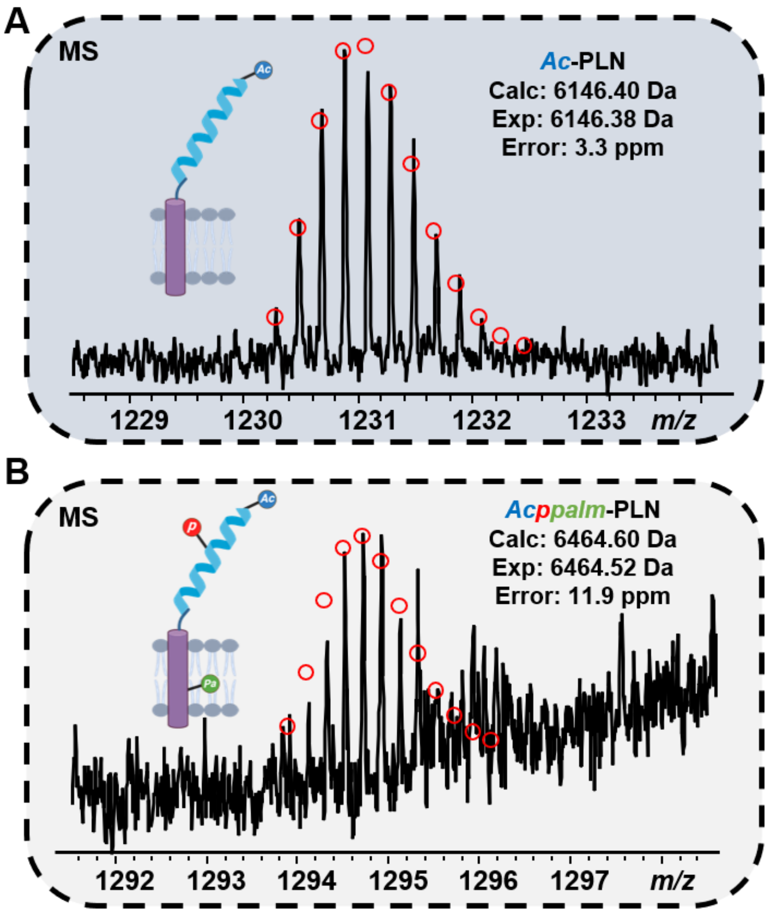


**SI Figure 4. Low abundant PLN proteoforms. (A)** MS of acetylated non-phosphorylated PLN proteoform. **(B)** MS of palmitoylated mono-phosphorylated PLN proteoform. Data collected using Impact II. Theoretical isotopic fits are shown in red circles. Monoisotopic masses are reported.

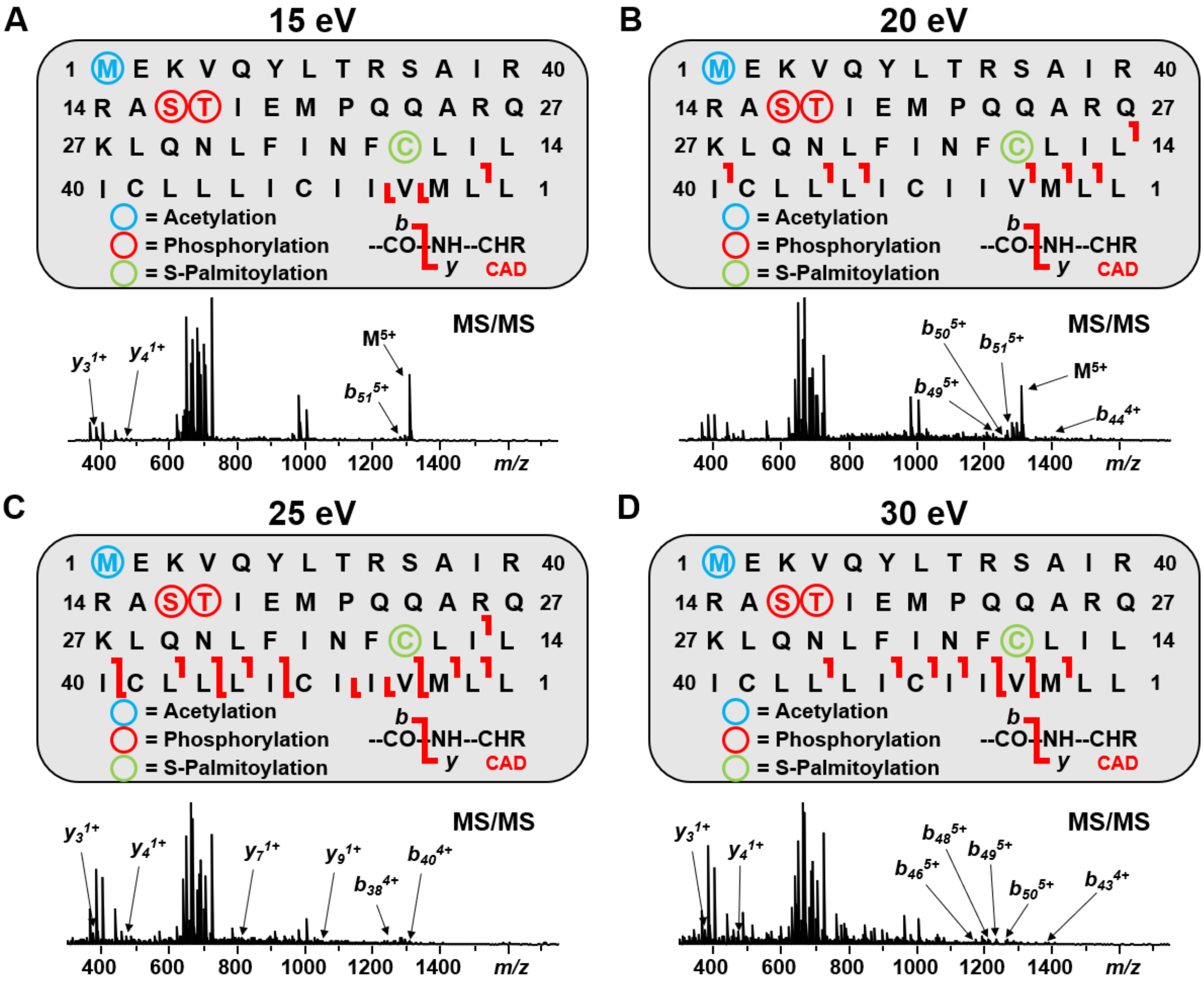


E

| Collision Energy (eV) | # of *b*-ions | # of *y*-ions | Bond Cleavage (%) |
|---|---|---|---|
| 15 | 1 | 2 | 5.77 |
| 20 | 7 | 0 | 13.46 |
| 25 | 9 | 6 | 21.15 |
| 30 | 7 | 2 | 13.46 |

**SI Figure 5. Online RPLC-MS/MS analysis of individual collision energies for the palmitoylated bis-phosphorylated PLN proteoform. (A-D)** Sequence tables report individual fragmentation coverage of PLN proteoforms across four different collisional energies: **(A)** 15 eV, **(B)** 20 eV, **(C)** 25 eV, and **(D)** 30 eV. Representative MS/MS spectra are displayed underneath their respective sequence table. Select fragments are annotated for each spectrum. **(E)** Table summarizing the bond cleavage percentage and number of *b*/*y*-ions produced during fragmentation with each collision energy.

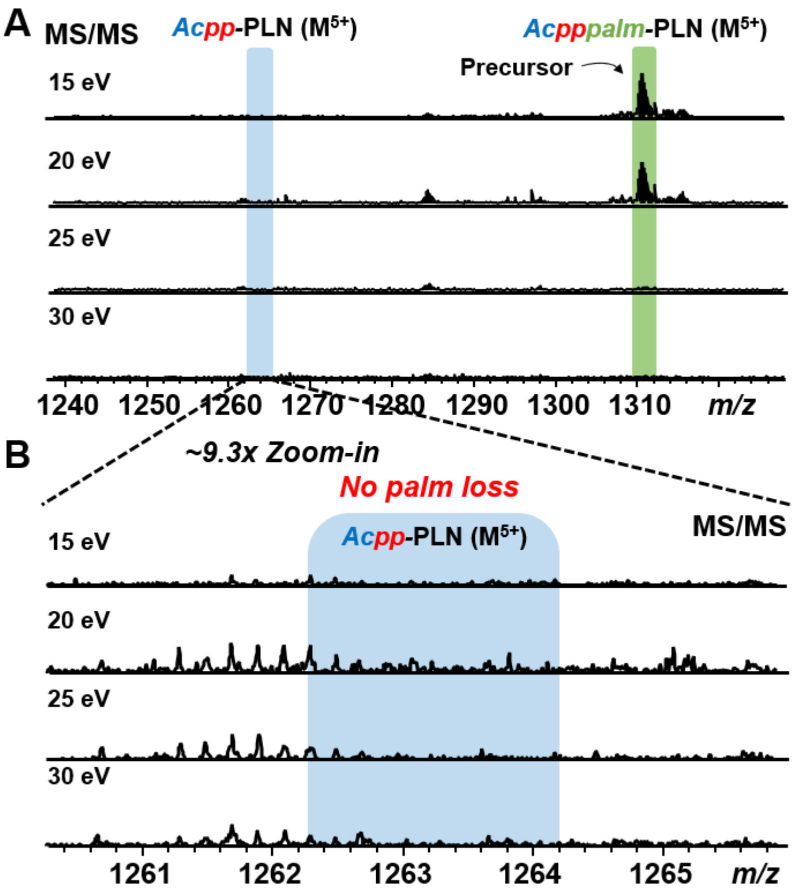


**SI Figure 6. Effect of CAD energy on palmitoylated bis-phosphorylated PLN proteoform. (A)** MS/MS spectra of palmitoylated bis-phosphorylated precursor upon CAD with increasing energy (15, 20, 25, and 30 eV). **(B)** Zoom-in on the *m/z* region where bis-phosphorylated PLN is found. Lack of presence of bis-phosphorylated PLN demonstrates minimal palmitoylation removal upon fragmentation.

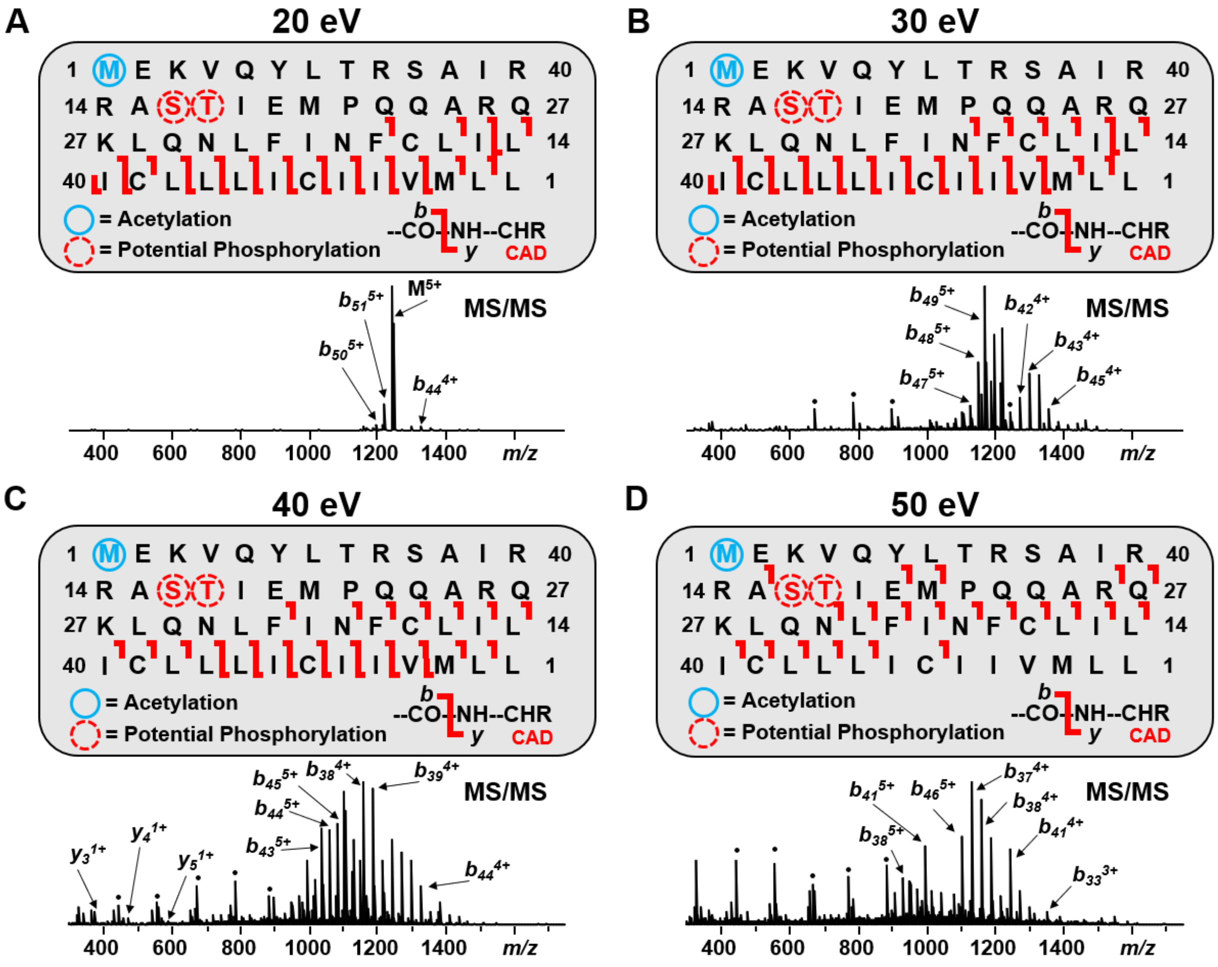


E

| Collision Energy (eV) | # of *b*-ions | # of *y*-ions | Bond Cleavage (%) |
|---|---|---|---|
| 20 | 16 | 11 | 30.76 |
| 30 | 18 | 12 | 34.62 |
| 40 | 19 | 7 | 36.54 |
| 50 | 21 | 0 | 40.38 |

**SI Figure 7. Online RPLC-MS/MS analysis of individual collision energies for the mono-phosphorylated PLN proteoform. (A-D)** Sequence tables report individual fragmentation coverage of PLN proteoforms across four different collisional energies: **(A)** 20 eV, **(B)** 30 eV, **(C)** 40 eV, and **(D)** 50 eV. Representative MS/MS spectra are displayed underneath their respective sequence table. Select fragments are annotated for each spectrum. Circles represent validated internal fragment ions corresponding to the mono-phosphorylated proteoform. Internal fragment ions were not included in bond cleavage percentage. **(E)** Table summarizing the bond cleavage percentage and number of *b*/*y*-ions produced during fragmentation with each collision energy.

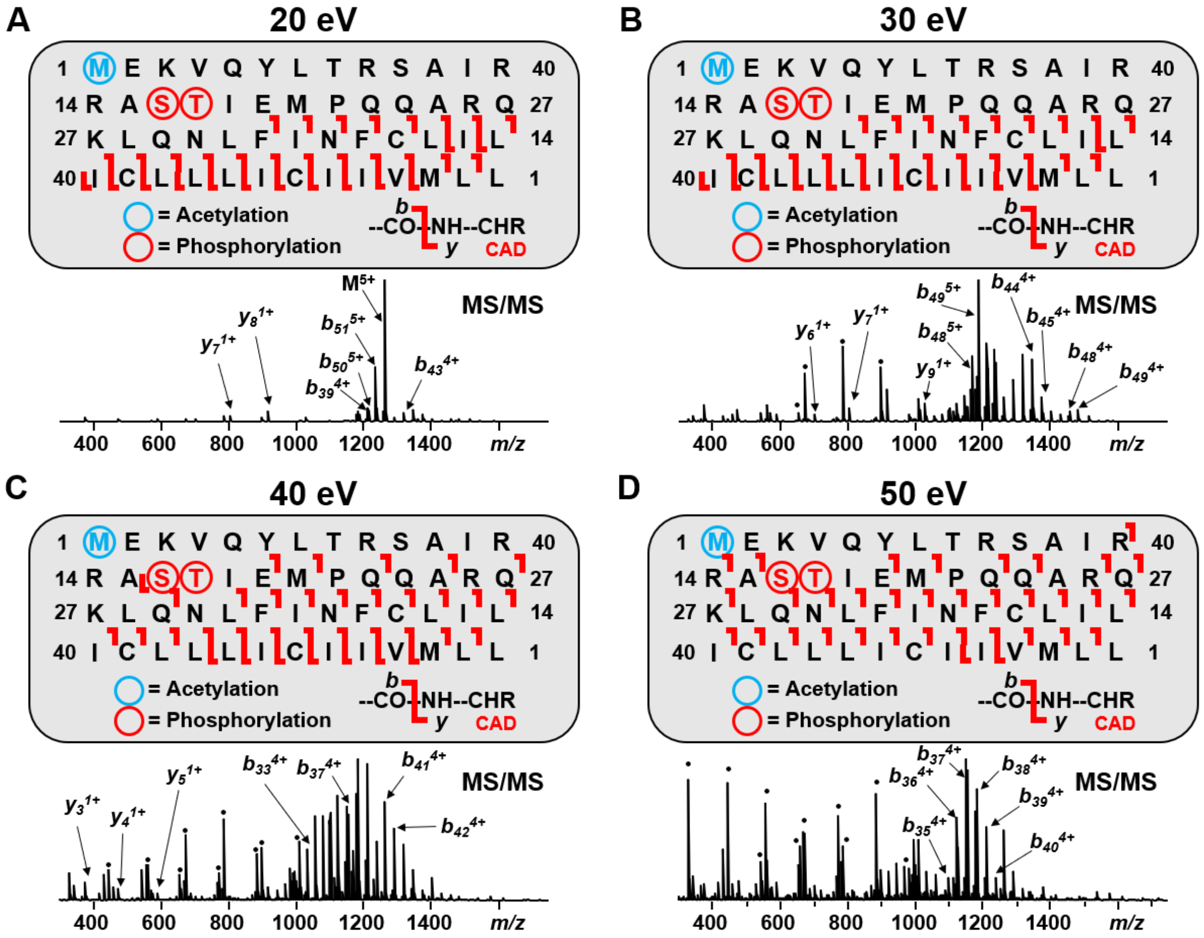


E

| Collision Energy (eV) | # of *b*-ions | # of *y*-ions | Bond Cleavage (%) |
|---|---|---|---|
| 20 | 20 | 13 | 38.46 |
| 30 | 21 | 12 | 40.38 |
| 40 | 27 | 8 | 53.85 |
| 50 | 33 | 2 | 63.46 |

**SI Figure 8. Online RPLC-MS/MS analysis of individual collision energies for the bis-phosphorylated PLN proteoform. (A-D)** Sequence tables report individual fragmentation coverage of PLN proteoforms across four different collisional energies: **(A)** 20 eV, **(B)** 30 eV, **(C)** 40 eV, and **(D)** 50 eV. Representative MS/MS spectra are displayed underneath their respective sequence table. Select fragments are annotated for each spectrum. Circles represent validated internal fragment ions corresponding to the bis-phosphorylated proteoform. Internal fragment ions were not included in bond cleavage percentage. **(E)** Table summarizing the bond cleavage percentage and number of *b*/*y*-ions produced during fragmentation with each collision energy.

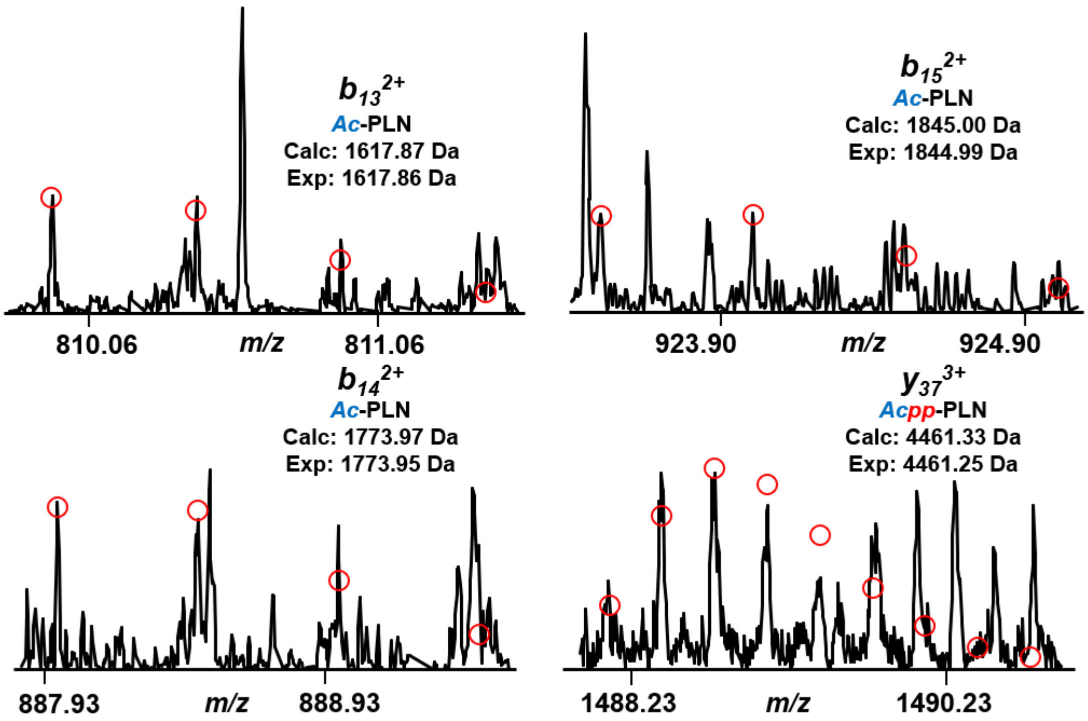


**SI Figure 9. Fragment ions supporting phosphorylation localization.** Spectra of PLN fragment ions ($b_{13}^{2+}$, $b_{14}^{2+}$, $b_{15}^{2+}$, $y_{37}^{3+}$) from online LC-MS/MS data that, in conjunction with FTICR data, support localization to serine-16 and threonine-17 rather than serine-10 or threonine-8.
